# Mutualistic cross-feeding in microbial systems generates bistability via an Allee effect

**DOI:** 10.1101/2019.12.23.886622

**Authors:** Stefan Vet, Lendert Gelens, Didier Gonze

## Abstract

Mutualistic interactions are characterized by the positive influence that two species exert on each other. Such mutualism can lead to bistability. Depending on the initial population size species will either survive or go extinct. Various phenomenological models have been suggested to describe bistability in mutualistic systems. However, these models do not account for interaction mediators such as nutrients. In contrast, nutrient-explicit models do not provide an intuitive understanding of what causes bistability. Here, we reduce a theoretical nutrient-explicit model of two mutualistic cross-feeders in a chemostat, uncovering an explicit relation to a growth model with an Allee effect. We show that the dilution rate in the chemostat leads to bistability by turning a weak Allee effect into a strong Allee effect. This happens as long as there is more production than consumption of cross-fed nutrients. Thanks to the explicit relationship of the reduced model with the underlying experimental parameters, these results allow to predict the biological conditions that sustain or prevent the survival of mutualistic species.

## Introduction

Microbes play a fundamental role in different ecosystems on Earth. For example, they provide nutrients for plants in the rhizosphere via a symbiotic relationship^1^, contribute to the formation of planktonic communities in the ocean^2,3^ and are used in the treatment of wastewater^4^. Even the human body is home to large ecosystems of microbial species, called microbiota, contributing to our health by providing essential nutrients and protecting us against potential threats or harmful microbial species^5^.

To understand the dynamics of microbial ecosystems, the growth of microbes can be studied *in vitro* under well controlled environmental conditions^6^. This way, microbes also provide convenient model systems to study general ecological interactions^7^. A particularly suited laboratory device to experimentally study microbial growth is the chemostat^8^. Such a bioreactor allows to grow microbes in a chemically constant environment and to explictly monitor the consumption of metabolites. A chemostat consists of a well-mixed growth tank with a continuous inflow of nutrients and an outflow of the suspension with microbes and nutrients. It is a simplification of natural systems as the inflow and outflow occur at the same rate and the suspension is well-mixed so that spatial effects are ignored. Nevertheless, it constitutes an appropriate tool to probe the behavior of natural systems, which are typically open environments with a flux of energy. For instance, it has been shown that the human intestines can, to some extent, be modeled by chemostat equations^9^, which are particularly suitable to assess the correlation between perturbed microbiomes (dysbiosis) and diseases. Experimental as well as theoretical studies involving a chemostat thus provide an appropriate framework to predict behavior related to microbial interactions such as competition and mutualism in a natural environment.

Although mutualism is thought to be less common than competition in microbial ecosystems because it tends to destabilize the community^10^, mutualism can arise via bi-directional cross-feeding of metabolites^11^. It has been shown that microbial diversity is promoted by cross-feeding, which can prevent competitive exclusion^12^. Furthermore, cross-feeding can be essential for different functions. For example, in the intestines metabolites are broken down in smaller components by some species for their consumption by other species^11^. This is necessary for the formation of health-promoting short-chain fatty acids^13–15^. Mutual cross-feeding has also been shown to reduce the energetic cost by dividing the labor for the utilization of metabolic pathways, for example for amino acid synthesis^16^.

Besides the apparent benefits of mutualism, there is a downside: the interdependency increases the possibility of a collapse of the system due to a density threshold for survival, which has been observed experimentally^17,18^. Mutualistic species are usually part of a larger ecosystem. Therefore, when mutualists are decreased under the survival threshold, for example in response to antibiotics, the entire ecosystem can be destabilized^10^. Such critical effects on ecological interactions are often not well characterized^19^. Therefore, in order to predict when such disruptions occur and, if needed, to intervene to prevent the collapse of the community, a deeper understanding of mutualistic interactions and of the occurrence of thresholds in microbial communities is necessary.

A density threshold between two different states of the system, in this case survival and extinction, is related to the concept of bistability. Different models have shown that mutualism can cause bistability via the interdependence between the species^20–23^. In some of these models the species are competitive as well as mutualistic, e.g. when mutualists become competitors for a resource or for available space at high density^24–26^. One study showed that the type of interaction could be modulated by varying the resource concentration^17^.

Mutualism has also been shown to create an Allee effect^27–29^. The presence of an Allee effect means that the individual growth rate reaches a maximum at an intermediate density. At low population densities, the microbial fitness thus benefits from an increased density. This effect arises in many cooperative systems and is in contrast with the prediction of the classical logistic growth which predicts that an increased population density limits the growth^30^. There is a distinction between a weak and a strong Allee effect^1^. Whereas a weak Allee effect leads to a single stable state (the species always survives), a strong Allee effect is characterized by bistability, whereby a density threshold for survival is present. A currently unresolved problem is that phenomenological models where interaction mediators, like nutrients, are neglected can behave differently than models where these are explicitly incorporated^31^. One approach, based on Mac-Arthur’s consumer-resource models, describes the occurrence of bistability by the saturation of mutualism at high densities^32^. Nevertheless, it remains unclear how the occurrence of bistability in a mutualistic cross-feeding community is related to nutrient concentrations and to their consumption and production kinetics. This is essential to quantify the effects of prebiotics or biological parameters on the survival of the species.

In this theoretical work, we use a nutrient-explicit model for the growth of mutualistic species in a chemostat reactor and show how biological parameters of this system are related to an Allee effect. This allows to predict when bistability is created and to estimate the density threshold for survival. Nutrient-explicit models of a single species in the chemostat can be reduced to the logistic growth equation^8,33,34^. Using a similar approach, we reduce chemostat equations of a mutualistic system to an appropriate mechanistic model which only involves the species densities. This allows to relate the obtained equations to a generic growth model with an Allee effect. By establishing this analogy, we show that mutualism causes a weak Allee effect, which can be turned into a strong Allee effect under the influence of the dilution in the chemostat. Critical chemostat parameters such as the dilution rate and the influx of nutrients thus allow to manipulate the strength of the Allee effect and therefore of the survival threshold. As a consequence, it is possible to switch between regions of bistability, monostable survival, or monostable extinction. We also show that the production of cross-feeding nutrients needs to be larger than the consumption for an Allee effect to exist. This explicit relationship between experimental parameters and the Allee effect provides a way to bridge the gap between biological experiments and theoretical models.

## Results

### Bistability in mutualistic systems creates a survival threshold

We study a theoretical system of two mutualistic species with densities *ρ*_1_ and *ρ*_2_. Mutualism is mediated by cross-feeding: each species consumes a nutrient, with resp. concentrations *P*_1_ and *P*_2_, produced by the other species (Fig 1A). We also assume that each species requires an additional nutrient which we refer to as the substrate, with resp. concentrations *S*_1_ and *S*_2_. The substrate consumption is necessary to avoid a violation of biomass conservation when production of the cross-fed nutrients is higher than the consumption. The mutualistic relationship creates a positive feedback loop which can lead to bistability. This phenomenon becomes apparent if we simulate the behavior of the species when cultured in a chemostat reactor (Fig 1). A chemostat consists of a well-mixed growth vessel with an inflow of nutrients, at concentrations 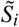 and 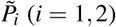, and an outflow of the suspension, which occurs with a dilution rate *d*. The consumption of nutrients is considered proportional to the growth of the species, which corresponds to the conservation of biomass when *d* = 0 (Supplementary Material S1). In the same way, we also assume the production of nutrients to be proportional to the growth of the species. The equations that describe the mutualistic chemostat system, with variables and parameters as defined in Table 1, are then:

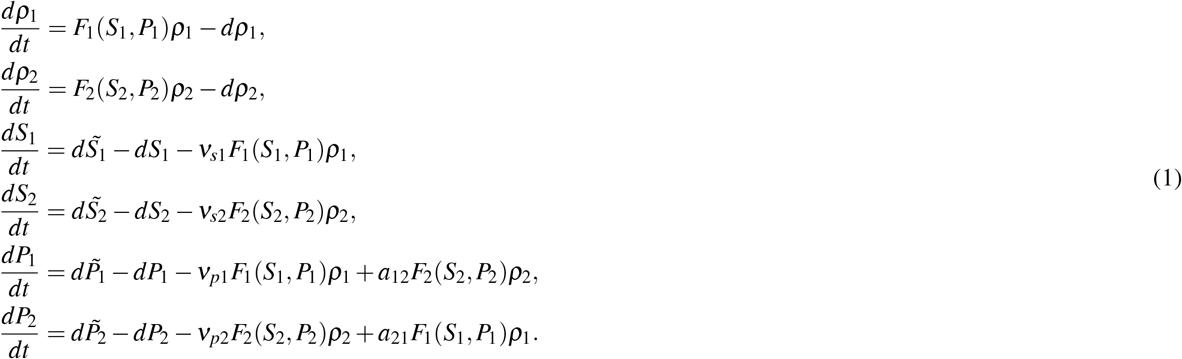

**Table 1.** Definition of the variables and parameters of nutrient-explicit chemostat equations of the mutualistic cross-feeding system, Eq (1), for *i* = 1,2 and *i* ≠ *j*.

| Variable | Biological meaning |
| --- | --- |
| $\rho_i$ | microbial density of species $i$ |
| $S_i$ | substrate of species $i$ (concentration) |
| $P_i$ | cross-fed nutrient, produced by species $i$ (concentration) |
| Parameter | Biological meaning |
| $d$ | dilution rate |
| $\tilde{S}_i$ | inflow concentration of $S_i$ |
| $\tilde{P}_i$ | inflow concentration of $P_i$ |
| $v_{si}$ | consumption rate of $S_i$ by species $i$ |
| $v_{pi}$ | consumption rate of $P_i$ by species $i$ |
| $a_{ij}$ | production rate of $P_i$ by species $j$ |
| $\mu_i$ | Maximal growth rate of species $i$ |
| $K_{si}$ | Monod constant for $S_i$ (consumption by species $i$) |
| $K_{pi}$ | Monod constant for $P_i$ (consumption by species $i$) |

**Figure 1.**
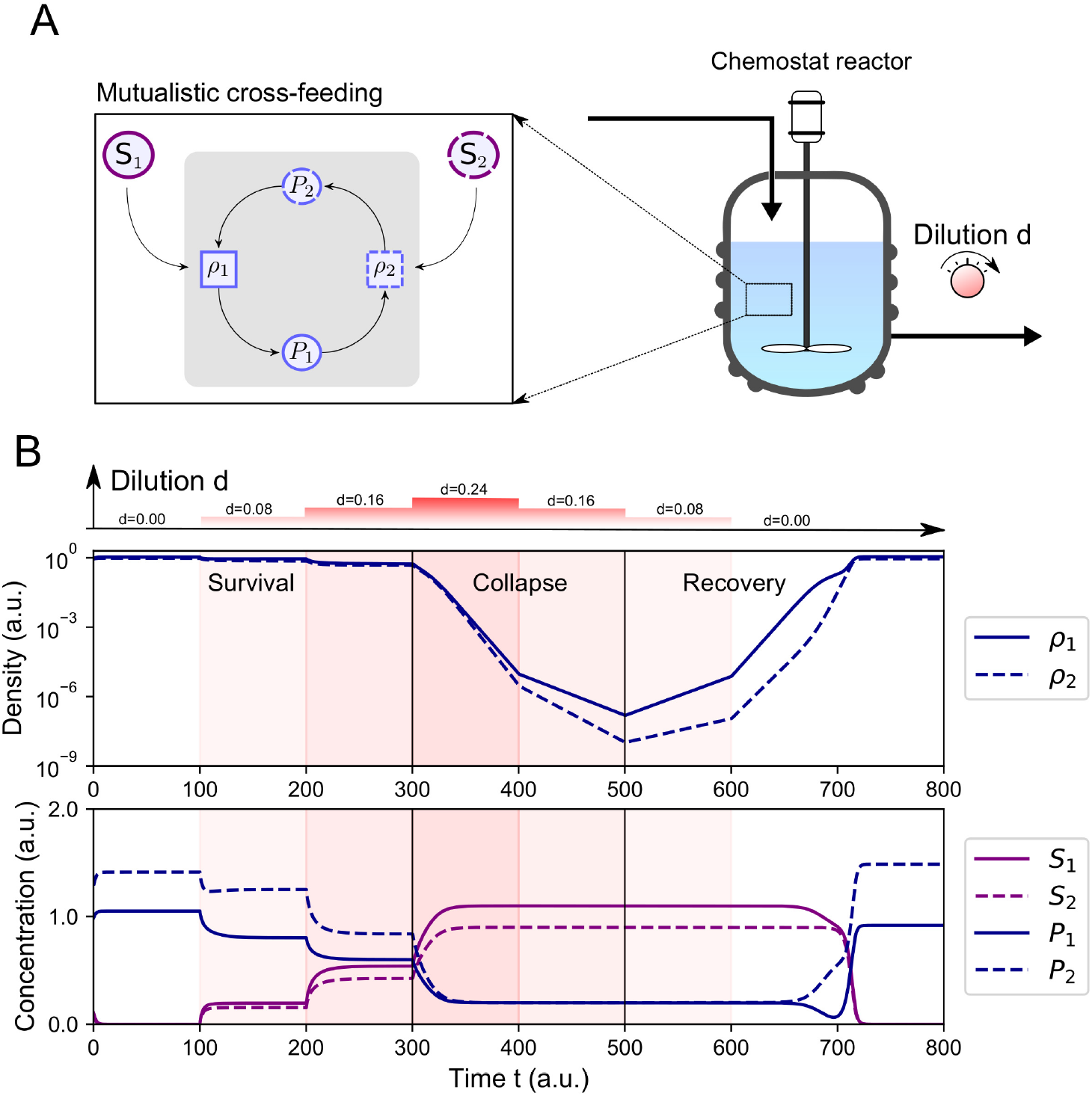
Bistability in mutualistic systems creates a survival threshold. (A) The microbial system we study consists of two species, characterized by their densities *ρ*_1_ and *ρ*_2_. Besides consumption of a substrate, resp. *S*_1_ and *S*_2_, the species engage in a mutualistic cross-feeding relationship via the production of nutrients resp. *P*_1_ and *P*_2_, which are consumed by the other species. We theoretically investigate the growth of these species in a chemostat reactor. This is a well-mixed growth vessel with an inflow of nutrients with concentrations 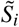, 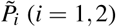 and an outflow of the suspension, occuring at an equal rate: the dilution rate *d*. These parameters can be adjusted to control or manipulate the dynamics. (B) The behavior of the system is simulated with Eqs (1) using the growth rate Eq (2), where we increase the dilution gradually from *d* = 0 to *d* = 0.24 with steps of 0.08. For clarity, we used a logarithmic scale for the species density. The equilibrium densities of the species decrease with the dilution. For *d* = 0.24, a threshold is reached and the species will be washed-out if the dilution remains unchanged. Trying to prevent wash-out by decreasing the dilution to its previous rate does not lead to the recovery of the initial abundances of the species, instead the dilution needs to be further decreased to *d* = 0.08 the make the population growing again. This is a phenomenon called hysteresis: the system has memory of the previous state. It is a consequence of bistability between survival and extinction at intermediate dilution, so that a density threshold for survival exists. (Parameters values: *µ*_1_ = 2, *µ*_2_ = 2, *K*_*s*1_ = 2, *K*_*s*2_ = 2, *K*_*p*1_ = 1, *K*_*p*2_ = 1, 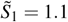, 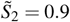, 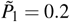, 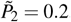, *ν*_*s*1_ = 1, *ν*_*s*2_ = 1, *ν*_*p*1_ = 1, *ν*_*p*2_ = 1, *a*_1_ = 2, *a*_2_ = 2)

We focus on obligate cross-feeding. Therefore, we assume obligatory dependence of the nutrients which means that the growth rate of the species (*F*_*i*_(*S*_*i*_, *P*_*i*_), with *i, j* = 1,2) needs to satisfy the following condition: *F*_*i*_(0, *P*_*i*_) = *F*_*i*_(*S*_*i*_, 0) = 0. An example of such a growth rate is the following extension of the Monod function for two nutrients^35^:

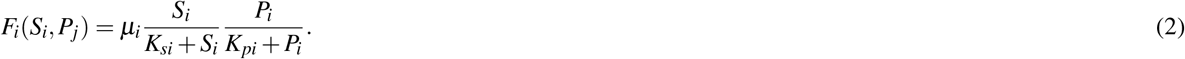

To gain some intuition about the system, we can simulate its behavior and consider what happens if the dilution is varied (Fig 1B). The growth rate becomes constant when equilibrium substrate concentrations are reached. In this state, the growth rate of each species equals the dilution rate *d*, allowing to manipulate the equilibrium. Increasing the dilution by one step (*d* = 0.08), the equilibrium density of a species decreases accordingly. However, a survival threshold exists: if the density drops below some critical values the species are washed-out. This is an irreversible collapse of the system, in the sense that reversing the dilution rate does not immediately lead to a return of the surviving state. Instead the species density continuous to drop. Only by a sufficient decrease of the dilution survival is restored. This concept is often called hysteresis: the state of the system is dependent on its history. It is an effect related to bistability, meaning that there are two distinct equilibrium states for the same environmental parameters. In this case there is bistability between survival and extinction of the two species. Even though this simulation provides some intuition, it is not straightforward to predict how the different parameters of the system affect bistability. This is essential to determine when the threshold for survival is crossed. To provide an answer, we show how these parameters affect the dynamics by revealing a close connection to the widely studied growth equation with an Allee effect.

### Dilution allows to switch between a weak and a strong Allee effect

Mutualistic cross-feeding is a form of cooperative behavior and causes a positive feedback loop between the two species. As a consequence, at small densities an Allee effect is created. The per capita growth rate of a species, a proxy for their fitness, increases with the density at small densities and it reaches a maximum at an intermediate density. This is in contrast with the logistic growth, where the per capita growth rate is maximal at zero density. There is an important distinction between a weak Allee effect, associated to monostable dynamics, and a strong Allee effect, associated to bistability. Here, we show that the mutualistic cross-feeding creates a weak Allee effect and that increasing dilution in the chemostat is able to turn this into a strong Allee effect. Dilution can thus promote bistability: the two species coexist via cooperation or both die (community collapses). The separation between the two types of behavior is determined by a threshold of the population size.

Based on the reduction of nutrient-explicit equations for the growth of one species in the chemostat to the logistic equation^8,33,34^(see Supplementary Material, section S1), we can reduce the explicit chemostat equations to a two-variable mutualistic system (Fig 2A). The calculations are detailed in the Supplementary Material (section S2). In brief, this reduction relies on the following assumptions (*i* = 1,2 and *i* ≠ *j*):

1. Conservation of biomass for *t* » 1/*d*:

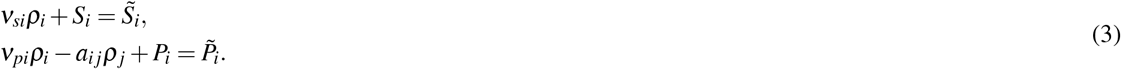 A Taylor approximation of the growth rate at low nutrient concentrations: 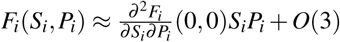

**Figure 2.**
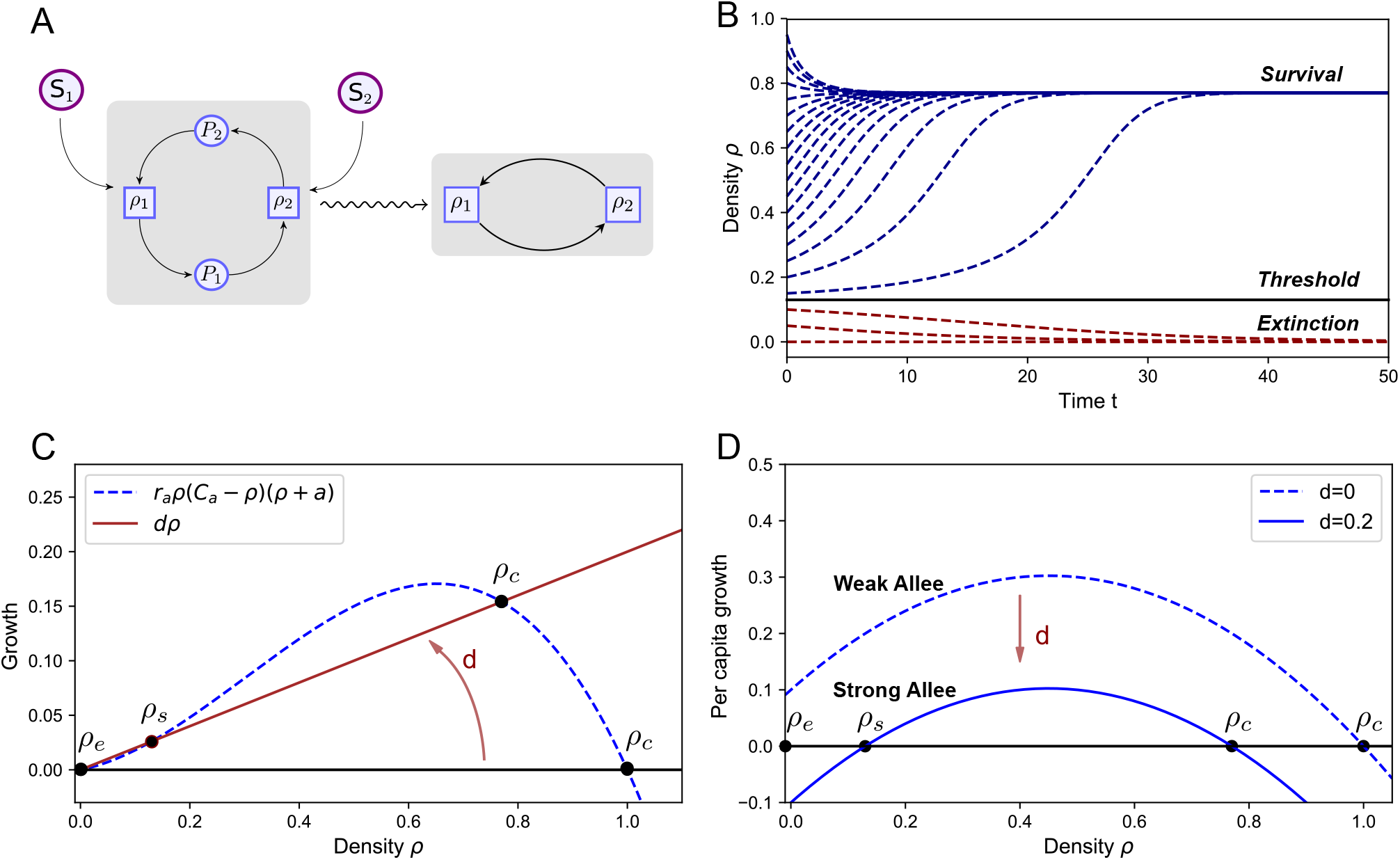
Dilution allows to switch between a weak and a strong Allee effect. (A) The nutrient-explicit chemostat equations, Eq (1), are reduced (wavy arrow) to equations which only involve the species densities, Eq (4), allowing to explain bistability in terms of an Allee effect. The mutualism causes an Allee effect via a positive feedback loop, which promotes bistability under the influence of the dilution rate. (B) Simulating the dynamics of Eq (5) shows the effect of bistability: depending on initial abundances, the species may coexist or become extinct. A threshold for survival separates the two outcomes. (C) The dilution rate is crucial for bistability. Without a dilution term, the system is monostable: there is only one stable equilibrium, corresponding to the survival of the species. Under the influence of the dilution, bistability occurs when there are three intersections of the growth and the dilution term: two stable fixed points corresponding to extinction and survival and an unstable steady state (saddle point) that separates both regions. (D) An Allee effect is present when the per capita growth increases with the density, which is the case for small densities and is the result of the positive feedback between the mutualistic species. For small dilution, the per capita growth does not become negative for small densities, corresponding to a weak Allee effect. When dilution is increased, the per capita growth rate becomes negative for small densities. This is a strong Allee effect and creates bistability between survival and extinction. Parameter values: *r*_*a*_ = 1, *a* = 0.1, *C*_*a*_ = 1, *d* = 0.2, see Supplementary Material, Table S7

The second assumption only holds in the case of obligatory dependence of the nutrients (*F*_*i*_(0, *P*_*i*_) = *F*_*i*_(*S*_*i*_, 0) = 0), such as the given example of the adapted Monod growth rate (Eq (2)).

Using a newly defined reduced set of parameters (see Table 2), we find that the two-species system can be described by the following equations:

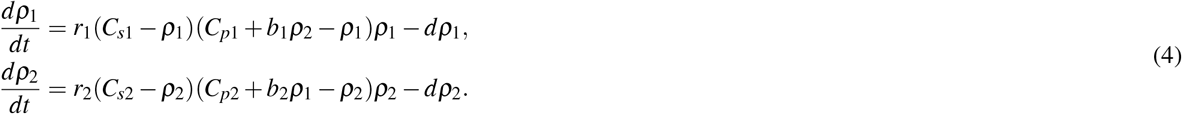

**Table 2.** Definition of the parameters for the reduced equations for the mutualistic cross-feeding system, Eq (4), as a function of chemostat parameters, for *i* = 1,2 and *i* ≠ *j*.

| Parameter | Biological meaning | Relation to chemostat system |
| --- | --- | --- |
| $d$ | dilution rate | $d$ |
| $r_i$ | growth rate | $v_{si} v_{pi} \frac{\partial^2 F_i}{\partial S_i \partial P_i}(0, 0)$ |
| $C_{si}$ | Carrying capacity on the limiting substrate | $\frac{\tilde{S}_i}{v_{si}}$ |
| $C_{pi}$ | Carrying capacity by a cross-fed nutrient | $\frac{\tilde{P}_i}{v_{pi}}$ |
| $b_i$ | Growth stimulation through cross-feeding | $\frac{a_{ij}}{v_{pi}}$ |

These equations can be interpreted as follows. Each species (with density *ρ*_*i*_) grows with a growth rate *r*_*i*_, which can be increased via mutualistic interactions (*b*_*i*_). The available nutrients increase with the other species population. As this is the case for both species, this results in positive feedback: the growth of each species is positively influenced by its own density. Such growth is limited by two carrying capacities, one related to the available substrate (*C*_*si*_) and one determined by the available cross-fed nutrients (*C*_*pi*_). The exact biological mechanism behind the limitation through these carrying capacities is not critical for the dynamics, which remain qualitatively similar even when species compete for the same substrate (see Supplementary Material, section S2). Additionally growth is further limited by the dilution in the chemostat (*d*).

In order to analyze the dynamics of this reduced equation (4), we first study the symmetric case where all parameters of both species are the same (e.g.: *r*_1_ = *r*_2_ = *r*). In this situation the dynamics maps to the subspace where the densities of the species are equal: *ρ*_1_ = *ρ*_2_ = *ρ*. The reduced mutualistic system is then found to be described by the following generic equation including cubic growth and an Allee effect^36^:

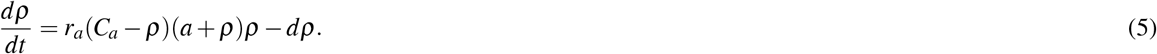

Here, the growth rate *r*_*a*_, carrying capacity *C*_*a*_, Allee parameter *a*, and loss term caused by the dilution (parameter *d*) are defined in table 3. The Allee effect is introduced by the factor (*a* + *ρ*) and is a consequence of the positive feedback between the species so that at low densities the per capita growth increases. Without dilution (*d* = 0) this equation takes the usual form of a growth equation with an Allee effect^36^. Based on the parameter *a*, the following distinction is made:

- *a* < 0: strong Allee effect (bistable)
- *a* > 0: weak Allee effect (monostable)

**Table 3.** Definition of the parameters for growth equation with Allee effect, Eq (5).

| Parameters | Biological meaning | Relation to reduced system |
| --- | --- | --- |
| $r_a$ | growth rate | $r(b - 1)$ |
| $C_a$ | carrying capacity | $C_s$ |
| $a$ | Allee parameter | $\frac{C_p}{b-1}$ |
| $d$ | dilution rate | $d$ |

A weak Allee effect corresponds to monostable growth towards an equilibrium, while a strong Allee effect is associated to bistability between survival and extinction so that a densitiy threshold for survival is generated (Fig 2B). As the Allee parameter is determined by the inflow of the cross-feeding nutrients and is therefore strictly positive, the mutualistic cross-feeding causes a weak Allee effect. The equilibria of the system are determined by the points where the growth is zero, which happens at *ρ* = 0 and *ρ* = *C*_*a*_ when *d* = 0 (Fig 2C). The extinction state *ρ* = 0 is unstable, while the coexisting state *ρ* = *C*_*a*_ is stable. For the chemostat case, the dilution is switched on (*d* > 0), the per capita growth is reduced by *d* and a new steady state may arise (Fig 2D). The new steady state is unstable forming the threshold for survival between the two stable equilibria (survival and extinction). This way, the dilution rate can turn a weak Allee effect into a strong Allee effect.

### Bifurcation analysis reveals regions of bistability

Before analyzing the biological consequences of the Allee effect in mutualistic interactions, it is useful to illustrate how the different parameters in Eq (5) affect the behavior. This can be done via a bifurcation analysis, showing the stability of the different equilibria as a function of the parameters.

The equilibria of the system correspond to the intersections of the growth term and the dilution term, where *dρ*/*dt* = 0, and are given by:

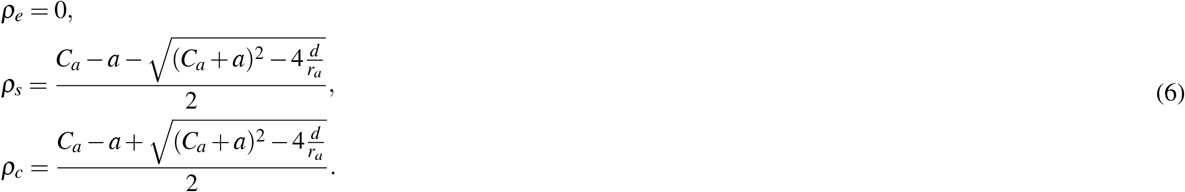

Here, *ρ*_*e*_ is the extinction state, *ρ*_*c*_ the coexisting state and *ρ*_*s*_ the separating state between the two equilibria in the case of bistability. The equilibria are determined geometrically by calculating how many times the per capita growth intersects with the dilution term (Fig 3 D,E). When there is only one intersection, the only stable equilibrium is *ρ*_*c*_ (*ρ*_*s*_ is negative and thus not physical). When there are two intersections, i.e. when *ρ*_*s*_ and *ρ*_*c*_ are both positive, the system is bistable: both the extinction state *ρ*_*e*_ and the coexisting state *ρ*_*c*_ are stable, and *ρ*_*s*_ is a so-called saddle point that separates the two stable equilibria. When the dilution rate is too large, there are no intersection and the net growth is always negative so that the only equilibrium is *ρ*_*e*_, corresponding to extinction of the species. In this manner, it is straightforward to calculate the values of the dilution rate where the behavior changes, leading to the following condition for bistability:

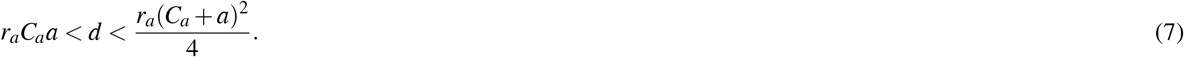

**Figure 3.**
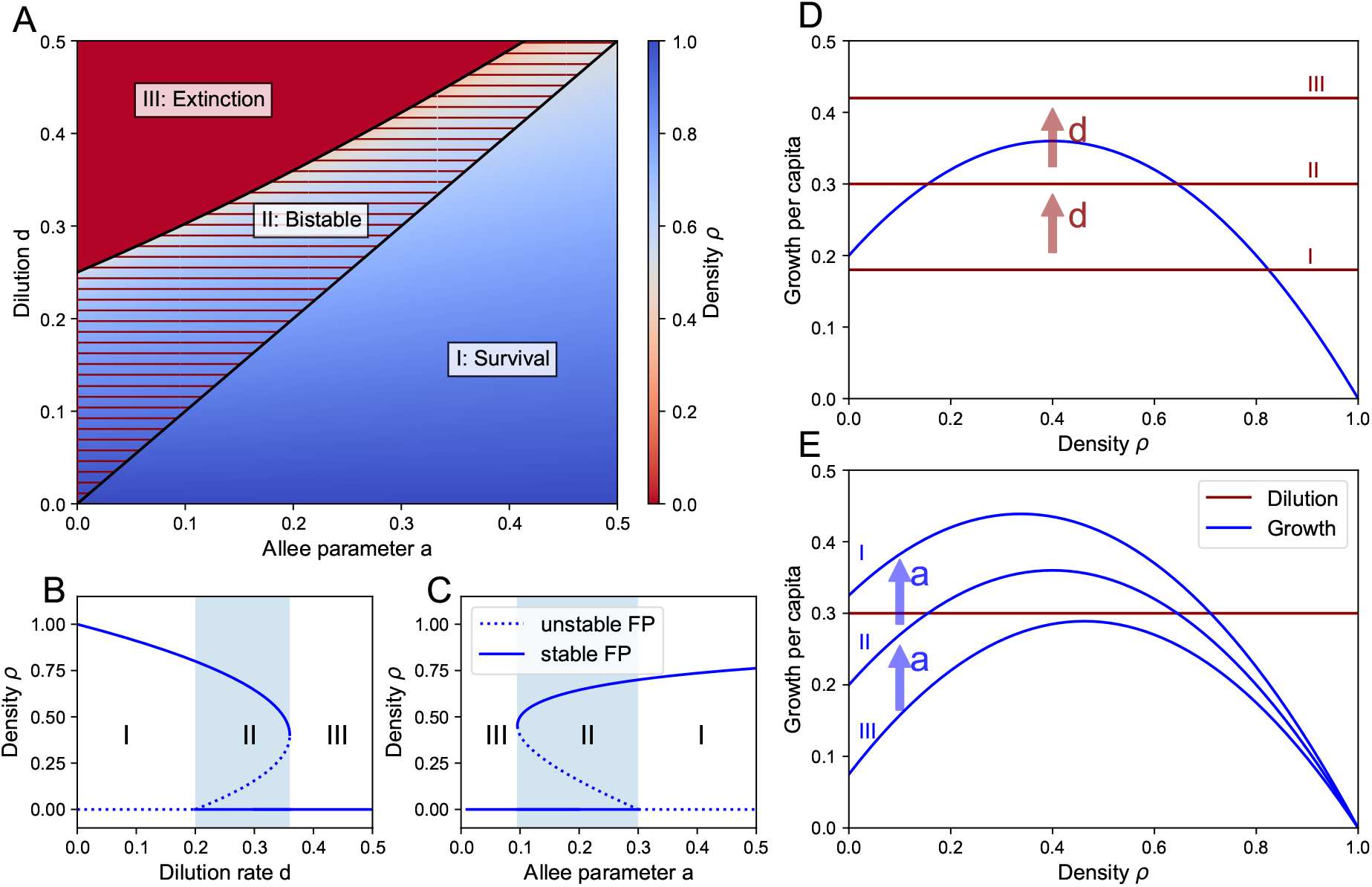
Bifurcation analysis reveals regions of bistability. (A) heatmap showing the equilibrium density *ρ* of Eq (5) as a function of the Allee parameter *a* and the dilution rate *d*. The different regimes correspond to survival (I), bistability between survival and extinction (II), and extinction (III). (B) Influence of the dilution *d*, for *a* = 0.2 kept constant. (C) Influence of the Allee parameter *a*, for *d* = 0.3 kept constant. (D) Visual interpretation of the equilibrium states, corresponding to the intersection of the per capita growth rate and the dilution, showing how increasing the dilution shifts regime I into regime II, into regime III. (E) The Allee parameter *a* has the opposite effect than the dilution *d*: increasing *a* allows for survival by shifting regime III into regime II into regime I. Parameter values: *r*_*a*_ = 1, *C*_*a*_ = 1.

For lower values of the dilution rate there is only survival and for higher values there is only extinction (Fig 3 A,B,D), thereby explaining the observed hysteresis in Fig 1. The same analysis shows that the Allee parameter *a* has a counter-acting effect on the dynamics in comparison to *d* (Fig 3 A,C,E): when the dilution rate is too large for survival, increasing *a* causes the per capita growth to intersect with the dilution, thereby allowing survival of the community. The complete behavior is summarized in Fig 3 A: for large values of *a* and low values of *d* there is always survival via cooperation (regime I), for intermediate values of *a* and *d* there is bistability (regime II) and for low values of *a* and large values of *d* there is always extinction of the community (regime III).

In order to check the validity of the used simplification of the growth rates, we simulated the original chemostat equations, Eqs (1), with extended Monod growth rates, Eq (2) (Fig 4). The used Taylor approximation of the Monod growth rates in the simplified model yields a higher value for the growth rate. Therefore, we expect extinction to occur at lower dilution than predicted with the simplified model (Fig 4A). We quantified the differences in the locations of the critical dilution rate *d*^∗^ where the saddle-node bifurcations (SN) occurs for both the full model and the reduced model (Fig 4 B). We performed the same calculation for different values of the Monod constants, whereby we multiplied the Monod constants by a factor *F* and the maximal growth rate *µ* by *F*^2^ so that the Taylor approximation of the growth rate remains the same. The same qualitative behavior was obtained when the Monod constants are of the order of the nutrients (F=1), but as expected a significant deviation was observed (Fig 4 C). Estimation of the error by considering the critical dilution rate *d*^∗^ for different values of *F*, shows that multiplying the Monod constants by a factor 10 yields a negligible error, i.e. a good quantitative match of the bifurcation diagrams was obtained with the explicit model and the reduced model (Fig 4 D).

**Figure 4.**
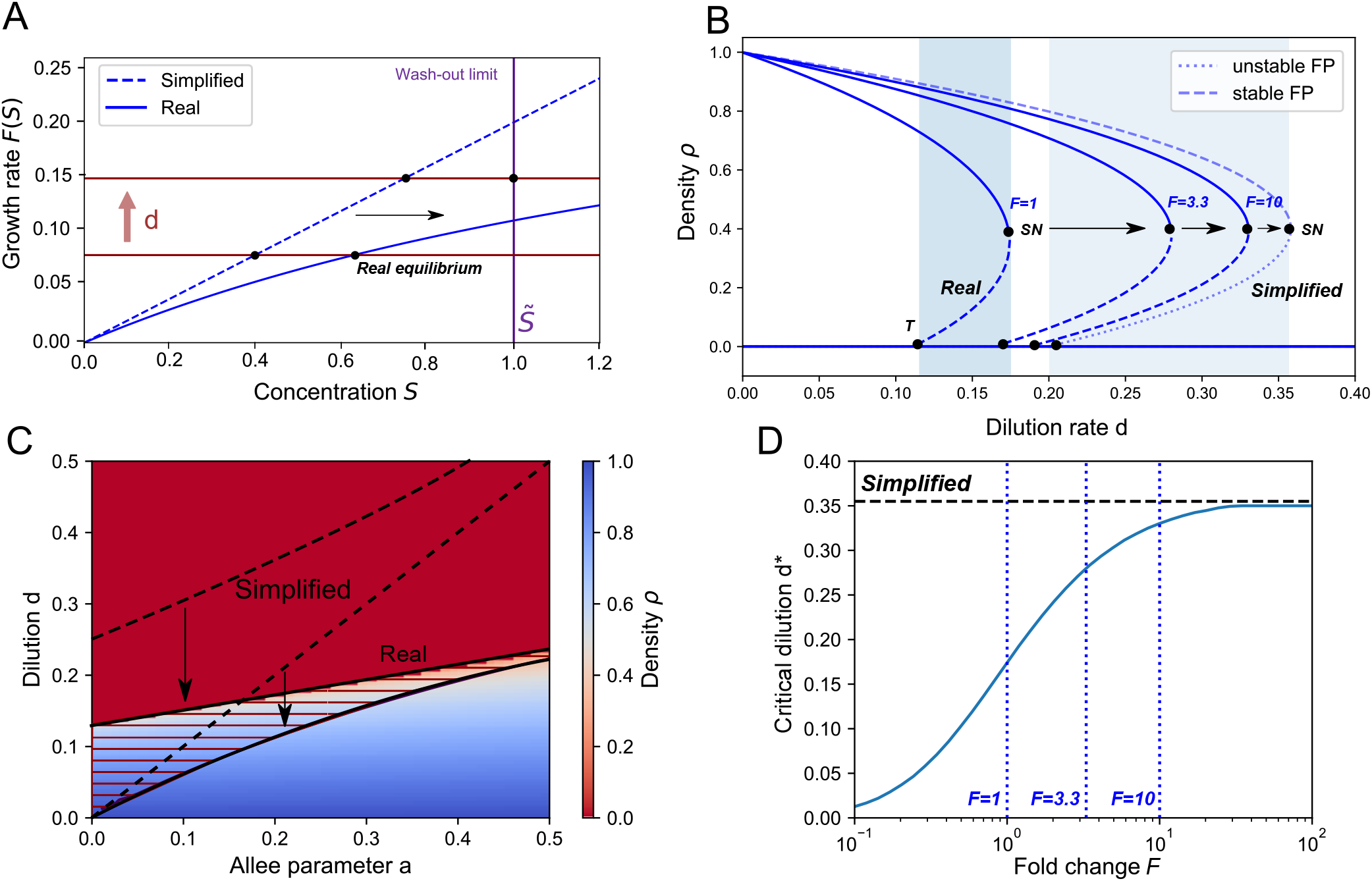
Original chemostat equations (1) confirm findings in the reduced equations (5). (A) The first-order Taylor approximation of the growth rate (Eq (2)) causes the wash-out limit to be reached for lower dilution rates. (B) By constructing the bifurcation diagram as a function of *d*, using chemostat equations (1) and the Monod-like growth rates, we can estimate the error by considering the point at which the saddle-node bifurcation (SN) occurs (T: transcritical bifurcation). A fold change *F* of the Monod constants is used to quantify the deviation from the simplified bifurcation curve. At *F* = 1 the error is large as *K*_*s*_ and *K*_*p*_ are of the order of *S* and *P* (Table S5). (C) Using *F* = 1, The calculated regions of survival, bistability and extinction in the reduced system (dashed curves) map on the simulated regions of the chemostat equations (solid curves) so that the qualitative behavior is conserved. (D) The error of the simplification is visualized by mapping the critical dilution rate at which the saddle node bifurcation (SN) occurs. For *F* = 1 the critical value of the dilution (*d*^∗^) is about 50% lower than the estimated value using the simplified model (Parameter values listed in Table S5).

### Bistability requires a sufficient production of cross-feeding nutrients

Bistability results from the mutualistic relationship between the two species. Having a dilution rate *d* in the correct parameter range is, by itself, not sufficient to guarantee bistability. Another necessary condition for bistability is obtained by considering the effect of the mutualistic interaction strength *b*, defined as the ratio of the production to the consumption of cross-fed nutrients 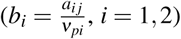. This parameter affects the growth rate *r*_*a*_ as well as the Allee parameter *a* (Table 3), so that its overall effect on the dynamics is not straightforward to predict intuitively. Therefore, we simulated the mutualistic system (Eq.(4)) with symmetric parameters, for different values of *b* and of dilution rate *d* (Fig 5 A). This analysis shows that the region of bistability is limited to *b* > 1. This means that bistability only occurs when the production of cross-fed nutrients is higher than the consumption. When *b <* 1, the production of cross-fed nutrients is insufficient. The growth is limited by these cross-fed nutrients, and the dynamics becomes similar as in the case of logistic growth, so that there is only one equilibrium state. The carrying capacity is then determined by the inflow of cross-feeding nutrients, via *C*_*p*_, and not by the inflow of substrate, via *C*_*s*_. Thus, for *b* < 1 there is no Allee effect.

**Figure 5.**
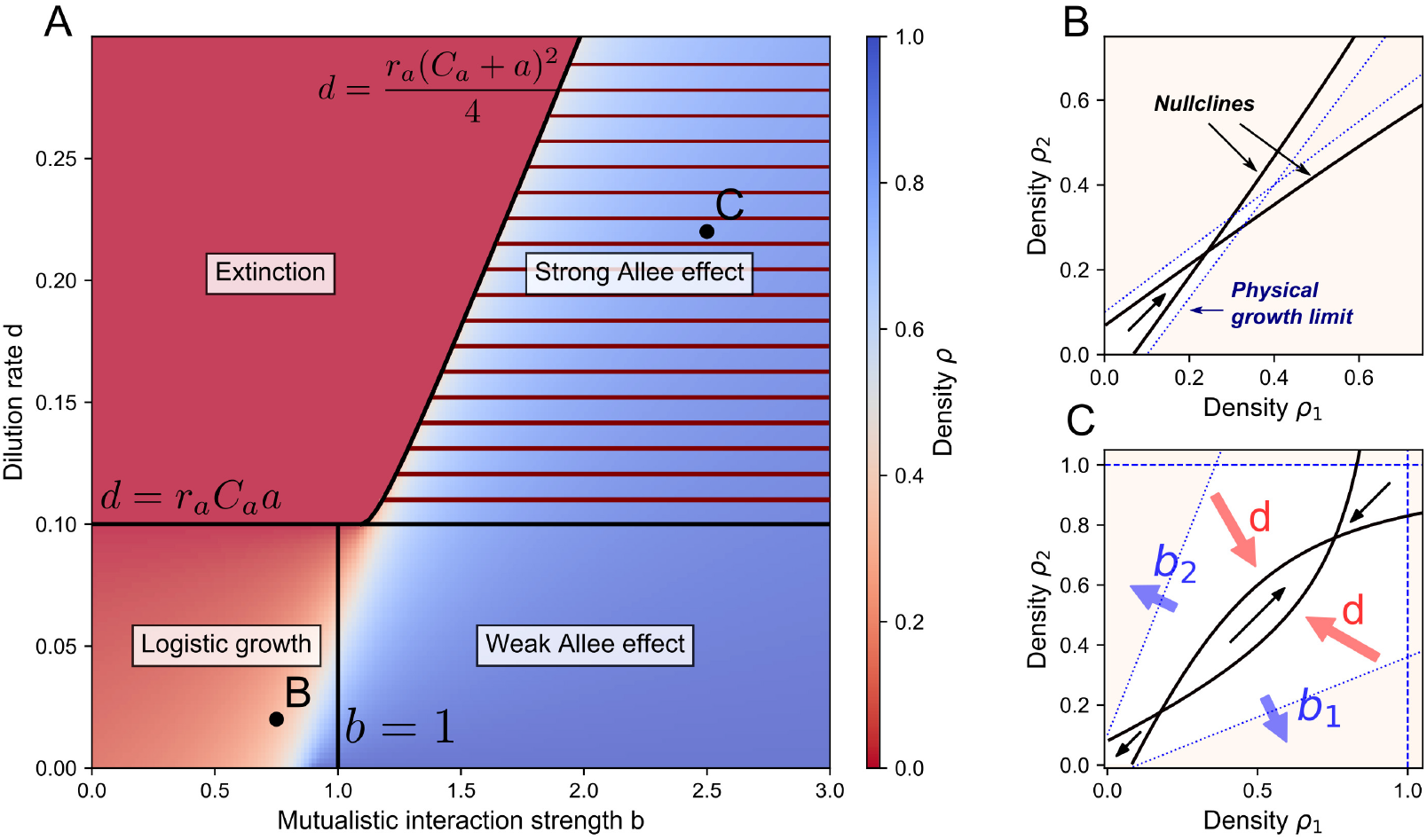
Bistability requires a sufficient production of cross-feeding nutrients. (A) Different regimes are distinguished as a function of the parameter values, using Eq (4). For *b <* 1, the growth is limited by the cross-feeding nutrient, leading to monostable dynamics similar to logistic growth. There is survival for small dilution (*d* < *r*_*a*_*C*_*a*_*a*) and extinction for large dilution. For *b >* 1, the growth only becomes limited by the substrate, allowing for bistability. There is a weak Allee effect, corresponding to monostable survival, for small dilution, a strong Allee effect for intermediate dilution 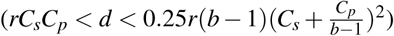 and monostable extinction for large dilution. (B) For *b <* 1, the growth is limited by the cross-feeding nutrients (Eq 9), so that the physical growth region is small and does not allow bistability. For small dilution the dynamics is similar to logistic growth as the population monotonically grows towards its equilibrium. (C) For *b >* 1, high population densities are possible, whereby the growth is limited by the substrate *S* (Eq 8). Increased values of *b* lead to a larger growth region, allowing for bistability when *b* > 1. Increasing the dilution rate *d* causes the nullclines to bend into hyperbola with the linear functions at *d* = 0 as asymptotes. Bistability is obtained when the hyperbolic nullclines intersect twice parameter values as listed in Table S6).

The difference between the logistic growth at *b* < 1 and the Allee effect at *b* > 1 is visually interpreted by representing the physical growth (i.e. feasible) regions in the phase plane for *b* < 1 (Fig 5 B) and for *b* > 1 (Fig 5 C). These feasible regions originate from the biomass conservation laws (Eq (3)), as the nutrient concentrations need to be positive. For each species, the limitation by the substrate causes the growth region to be bounded by the following function (*i* = 1,2):

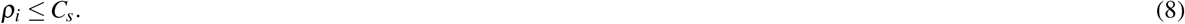

Similarly for the cross-feeding nutrients the following limitation functions are obtained (*i* = 1,2 and *i* ≠ *j*):

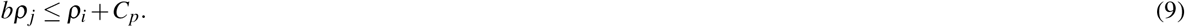

The interaction strength *b* determines the slope of Eq (9), so that for *b* < 1, the growth is entirely limited by Eq (9) and thus by the availability of cross-feeding nutrients (Fig 5 B). The species cannot grow sufficiently to become limited by the substrate. In contrast, for *b* > 1, the species can grow sufficiently. In this case, the feasible region is limited at high densities by the availability of the substrate, Eq (8) (Fig 5 C).

The overall influence of the parameters is understood by looking at the nullclines: the functions where *dρ*_*i*_/*dt* = 0 (*i* = 1,2). The intersections of the nullclines define the equilibria of the system. The nullclines are hyperbolic functions, with the limiting functions (Eq (8) and Eq (9)) as asymptotes. For bistability, the slopes need to be such that the nullclines can intersect twice (Fig 5 C), thereby forming a stable equilibrium corresponding to coexistence of the species and a saddle point which creates the threshold between coexistence and survival. This only occurs when *b* > 1, so that in this region we observe the same dynamics as described in last section: survival when the hyperbola intersect once at low dilution, bistability when the nullclines intersect twice for intermediate dilution 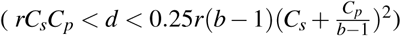 and extinction for high dilution (for the phase plane analysis, see Supplementary Material, Fig S3).

Bistability is created in the same way when asymmetric parameters are used. A similar analysis was performed for asymmetric values of the mutualistic interaction strength *b* (Supplementary Material, Fig S5 and Section S3), resulting in the general necessary condition for bistability: *b*_1_*b*_2_ > 1. Finally, the same result holds when the two mutualistic species compete for a substrate (Supplementary Material, Fig S4)). The competition reduces the physical growth region so that the species survive at lower densities, but the qualitative effect of the dilution rate and the mutualistic strength remains the same. Returning to the chemostat parameters, the obtained condition for bistability *b*_1_*b*_2_ > 1 corresponds to *a*_1_*a*_2_ > *ν*_*p*1_*ν*_*p*2_, which means that the overall production of cross-fed nutrients needs to be larger than their consumption.

## Discussion

Microbial interaction networks are characterized by multiple positive and negative interactions. Species enter in competition for limited resources, but they can also display mutualistic relationships through cross-feeding. Through mutualistic interactions, both species benefit of each others presence. This may be seen as a stabilizing factor. However, mutualism carries the seed of its own instability: under the influence of dilution bistability may occur, causing a critical density threshold for the species to survive. Once the abundances of the species drop below this threshold, the community eventually collapses and the species become extinct.

How biological parameters affect the survival threshold is often unclear. To provide an understanding of the effect of different parameters, we showed how nutrient-explicit equations for two mutualistic cross-feeding species can be reduced to a set of equations which only involve the densities of the species. These could then be related to a growth equation with an Allee effect, which can be analyzed to obtain a deeper understanding of the impact of the different biological parameters. We obtained quantitative results for the case where both species have symmetric parameter values, but showed that this framework still applies to the case of asymmetric parameter values. Our results showed that the overall production rate of cross-fed nutrients needs to be larger than the overall consumption rate to create an Allee effect. The production and consumption rates can experimentally be altered by making use of synthetic cross-feeding systems^17,37,38^, which can be designed to display bistability, or on the contrary, to prevent bistability and thereby the risk of an abrupt collapse of the community. We also showed that the dilution rate and the influx of nutrients are experimental parameters which can be tuned to manipulate the behavior. The effect of prebiotics can be simulated via the influx of nutrients (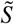 and 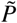), in order to determine the effect of these growth-promoting nutrients on the survival of the species. Furthermore, if antibiotics are used, then determining whether a survival threshold exists would be essential to predict the survival or extinction of the species.

Density thresholds for survival have previously been observed in mutualistic systems. For example, a survival threshold was found in a spatially cooperating microbial community^18^ and in a cross-feeding system which could switch to other interactions like competition and parasitism, depending on the availability of nutrients^17^. Besides cross-feeding, mutualism can also arise via the protection of another species towards antibiotics. In such a cross-protection system it was found that periodic dilution drives oscillatory dynamics, potentially leading to extinction if the survival threshold was crossed^39^. Recently, we were capable to fit a growth model on an experimental system of 3 species involving mutualistic cross-feeding in a batch reactor^40^, but as these species were shown to be facultative cross-feeders, bistability is not expected in this system. Nevertheless, the same fitting procedure could be used for obligate cross-feeding systems, in which case the theoretical predictions can be experimentally verified.

Another disadvantage of the existence of a survival threshold is that it creates susceptibility to cheaters^23^. A cheater is an individual of the species which does not cooperate in the creation of cross-feeding nutrients, thereby creating an energetic advantage over the cooperators and potentially increasing the risk of a collapse of the system^41^. On the other hand, by using a phenomenological model it has been shown that addition of a third species can create global stability of the coexistence state if it is a facultative mutualist, in which case there is no longer a survival threshold^42^.

Furthermore, when the spatial expansion of a species is considered, an Allee effect counteracts genetic drift of a species as it creates a pushed wave rather than a pulled wave corresponding to logistic growth^43,44^. This has been observed experimentally in a system of two cross-feeding species whereby the mutualistic strength was modulated by the inflow of nutrients^45^.

Our model involved the presence of a substrate due to biomass conservation, which limits the growth so that divergences are not possible. We modeled the substrate as a nutrient, but it can also be interpreted as the limited availability of space^26^. In fact, the obtained phase plane and the nullclines, describing the dynamics of the species, are observed in different mutualistic models^20–23,46^, so that we can state this is a general phenomenon. Therefore, our results give insights into necessary conditions for obligate mutualistic models: there needs to be an Allee effect as well as a limiting function. Lotka-Volterra models for mutualism do not incorporate these conditions^47^, as these involve the addition of fitness effects^31,48^. Therefore, our results hint at the use of nonlinear growth rates in generalized Lotka-Volterra models to study the survival of species in ecosystems involving obligate mutualism.

## Acknowledgements

This work was supported by the “Fonds National de la Recherche Scientifique” (FNRS, Belgium) (Project CDR J.0076.19).

## Author contributions statement

SV conceived the study, performed the analyses, and wrote the manuscript. LG and DG supervised the work and revised the manuscript. All authors gave final approval of the manuscript.

## Additional information

### Competing interests

The authors declare no competing interests.

