## Supplementary Material for "Mutualistic cross-feeding in microbial systems generates bistability via an Allee effect"

### Contents

**S1** Reduction of nutrient-explicit equations of a single species. **2**

**S2** Reduction of nutrient-explicit mutualistic cross-feeding equations **7**

**S3** Analysis for asymmetric parameter values **13**

References **14**

### S1 Reduction of nutrient-explicit equations of a single species.

#### Batch and chemostat reactors

The reduction of consumer-resource models requires to make the distinction between batch and chemostat reactors. We summarize the main differences here. Broadly speaking, a batch reactor corresponds to a closed environment, while a chemostat reactor corresponds to an open environment. This means that, in batch, the amount of biomass that can be produced is limited by the initial concentration of available nutrients, whereas in a chemostat there is a continuous flow of nutrients, allowing the microbial species density to reach a specific equilibrium state.

Both reactors are used to study the growth of bacteria *in vitro*. A batch reactor is a well-mixed, closed vessel with a suspension that is a mix of the substrate (concentration $S$) and of microbial species (density $\rho$). As there is only a limited amount of substrate available for the consumption, the species grows until the substrate is depleted. Therefore, the final population size is determined by the amount of species and substrate that are initially in the reactor, i.e. the initial concentrations.

A chemostat is an extension of the batch reactor with different dynamics as it is an open environment. There is a continuous inflow of nutrients with concentration $\tilde{S}$, and a continuous outflow of the suspension. The inflow and the outflow happen at the same rate: the dilution rate $d$. In the following we show that the nutrient-explicit equations for the growth of a microbial species in batch and in chemostat equations can be reduced to the logistic growth equation, which is a well-known result[1–3].

We consider a single species (density $\rho$) that consumes the substrate (concentration $S$), which it needs in order to grow. First, we consider the growth in a batch reactor, for which the equations can be reduced to a logistic growth equation. Then, we consider the growth in chemostat and show that a similar reduction can be done. Finally, we discuss the dynamics, whereby the difference between the two systems is essentially captured by the dilution rate.

#### Reduction of the batch equations

**Table S1.** Variables for the growth equations of a single species (Eqs (S4) and Eq (S3))

| Variables | Biological meaning | Dimension |
|---|---|---|
| $\rho$ | microbial density | arbitrary units (a.u.) |
| $S$ | substrate concentration | a.u. |

**Table S2.** Parameters of the chemostat model for a single species (Eqs (S4))

| Parameters | Biological meaning | Dimension | Value |
|---|---|---|---|
| $d$ | dilution rate | time$^{-1}$ | 0.2 |
| $\tilde{S}$ | inflow concentration of substrate $S$ | a.u. | 1 |
| $\mu$ | maximal growth rate | time$^{-1}$ | 1 |
| $K$ | half-saturation constant | a.u. | 1 |
| $\nu$ | consumption rate | dimensionless | 1 |
| $Y$ | yield: production of biomass per consumed substrate | dimensionless | $\frac{1}{\nu}$ |

If a species grows in a closed environment, corresponding with a batch reactor, then the growth of a species (density $\rho$) on a single resource (concentration $S$) is described by the following equations:

$$\begin{aligned} \frac{d\rho}{dt} &= F(S)\rho \\ \frac{dS}{dt} &= -\frac{F(S)\rho}{Y} \end{aligned} \tag{S1}$$

**Table S3.** Parameters of the logistic model for a single species (Eq (S3) and (S5))

| Parameters | Biological meaning | Dimension | Relation to chemostat system | Relation to batch system | Value |
|---|---|---|---|---|---|
| $d$ | dilution rate | time$^{-1}$ | $d > 0$ | $d = 0$ | 0.2 |
| $r$ | growth rate | time$^{-1}$ | $\frac{F'(0)}{Y}$ | $\frac{F'(0)}{Y}$ | 1 |
| $C$ | carrying capacity | a.u. | $Y\tilde{S}$ | $\rho(0)+YS(0)$ | 1 |

Here, $F(S)$ is a general growth rate function that depends on the substrate concentration, while $Y$ is the yield, i.e. the net growth per consumed substrate. It follows that this system obeys the following conservation equation:

$$\frac{d\rho}{dt} + Y\frac{dS}{dt} = 0$$

Hence, the quantity $\rho + YS$ is conserved and the substrate concentration $S$ depends linearly on the species density $\rho$:

$$S = \frac{\rho(0)+YS(0)-\rho}{Y}$$

It is assumed that the growth is limited by a low substrate concentration, meaning that $F(0) = 0$. Therefore, we can approximate the growth rate $F(S)$ to be linear in $S$, corresponding to the first-order term of the Taylor approximation:

$$F(S) \approx F(0) + F'(0)S + O(2)$$

A typical growth rate is the Monod function with $\mu$ the maximal growth rate and $K$ the half-saturation constant:

$$F(S) = \mu\frac{S}{K+S} \tag{S2}$$

As there is no growth when the substrate concentration is zero ($F(0) = 0$), the original set of differential equations simplifies to:

$$\frac{d\rho}{dt} = F'(0)\frac{\rho(0)+YS(0)-\rho}{Y}\rho$$

Redefining the parameters, the standard logistic equation with growth rate $r$ and carrying capacity $C$ is obtained:

$$\frac{d\rho}{dt} = r(C-\rho)\rho \tag{S3}$$

Notice that this restricts the physical values of $\rho$. The case $C - \rho < 0$ would correspond to negative values of $S$ and is therefore unrealistic. Therefore, this description is limited to the part of the phase space where $\rho < C$.

### Reduction of the chemostat equations

When microbial growth in a chemostat is considered, the equations are changed to allow an inflow of nutrients with concentration $\tilde{S}$, and an outflow of the suspension at the dilution rate $d$:

$$\begin{aligned}\frac{d\rho}{dt} &= F(S)\rho - d\rho,\\ \frac{dS}{dt} &= -\frac{F(S)\rho}{Y} + d\tilde{S} - dS.\end{aligned} \tag{S4}$$

The dilution rate $d$ determines the speed by which the substrate is added and by which the complete suspension is flown out of the system. As we will see, it is an essential parameter to determine the qualitative dynamics of the system. In a chemostat, in contrast to batch reactors, the total biomass $\rho + YS$ is initially not conserved. However, considering the same linear combination of variables, a slightly different but similar conservation law is obtained. The differential equation for $\rho+YS$ is:

$$\frac{d\rho}{dt}+Y\frac{dS}{dt}=d\tilde{S}-d(\rho+YS).$$

This has the following analytic solution:

$$\rho+YS=Y\tilde{S}+(\rho(0)+YS(0)-Y\tilde{S})e^{-dt}.$$

The initial concentrations thus play a role for $t<1/d$. For $t=0$, the same biomass conservation law as in batch is obtained. This is understood by the only time-dependent term, the exponential factor $e^{-dt}$, which decreases with a rate $d$, so that at $t\gg 1/d$, this term approaches zero. Thus, after an initial time $t\gg 1/d$, the following biomass conservation law is obtained:

$$\rho+YS=Y\tilde{S}$$

Therefore, the substrate concentration is given by $S=\tilde{S}-\frac{\rho}{Y}$. Substitution in Eq.(S4), yields a similar logistic equation as in batch, with altered parameters (Table S2) and with a loss term that is caused by the dilution rate:

$$\frac{d\rho}{dt}=r(C-\rho)\rho-d\rho. \tag{S5}$$

The effect of the dilution rate is straightforward: it reduces the carrying capacity to $C'=C-d/r$ and the per capita growth is reduced by $d$. In fact, by using $S(0)=\tilde{S}$, we see that for $d=0$ the equation reduces to the result for a batch reactor. Therefore, no dilution ($d=0$) corresponds with a batch reactor and a dilution larger than zero ($d>0$) corresponds with a chemostat reactor.

To summarize, in this reduction of a resource-explicit model we used the following approximations:

- The growth rate is linear at low substrate concentration so that: $F(S)\approx F'(0)S+O(2)$.
- By making use of the conservation of biomass, equations for the substrate can be removed.

### Dynamics of the logistic growth equation

We showed that the growth of a single species can be described by the logistic growth equation, which is a well-known result[1–3]. This model accounts for the limitation of the growth when a carrying capacity $C$ is reached, which is due to the depletion of the available nutrients in the environment. Here, we show how the dilution in the chemostat affect the dynamics of the logistic growth equation (Eq (S3)).

The relationship between the parameters of the chemostat model and the logistic equation is given in Table S3. When the microbial growth is simulated using different initial concentrations, the species always grows towards a specific density (Fig. S1B). When the species grows in batch ($d=0$) instead of in a chemostat, there is no dilution and the equilibrium concentration is the carrying capacity. In an open environment like the chemostat, dilution causes a lower effective carrying capacity $C'=C-d/r$. This is understood by calculating the steady states of Eq (S3). These are the equilibrium densities and correspond to the zero-growth solutions, $\frac{d\rho}{dt}=0$ so that these are visualized by the intersection of the growth term and the dilution term (Fig. S1C), resulting in the following states:

$$\begin{aligned}\rho_e&=0,\\ \rho_c&=C-\frac{d}{r}.\end{aligned} \tag{S6}$$

As long as $\rho_c>0$, this equilibrium is the only stable steady state, while $\rho_e$ is unstable. An equivalent, but more convenient way to visualize the effect of the dilution rate is by considering the per capita growth $\frac{1}{\rho}\frac{\rho}{dt}$: the net growth per individual, which can be interpreted as the fitness of the individual. This shows that the net per capita growth is reduced by $d$ (Fig. S1D), but there is no qualitative change in the dynamics: the stability of the equilibria remains the same as long as $d<rC$.

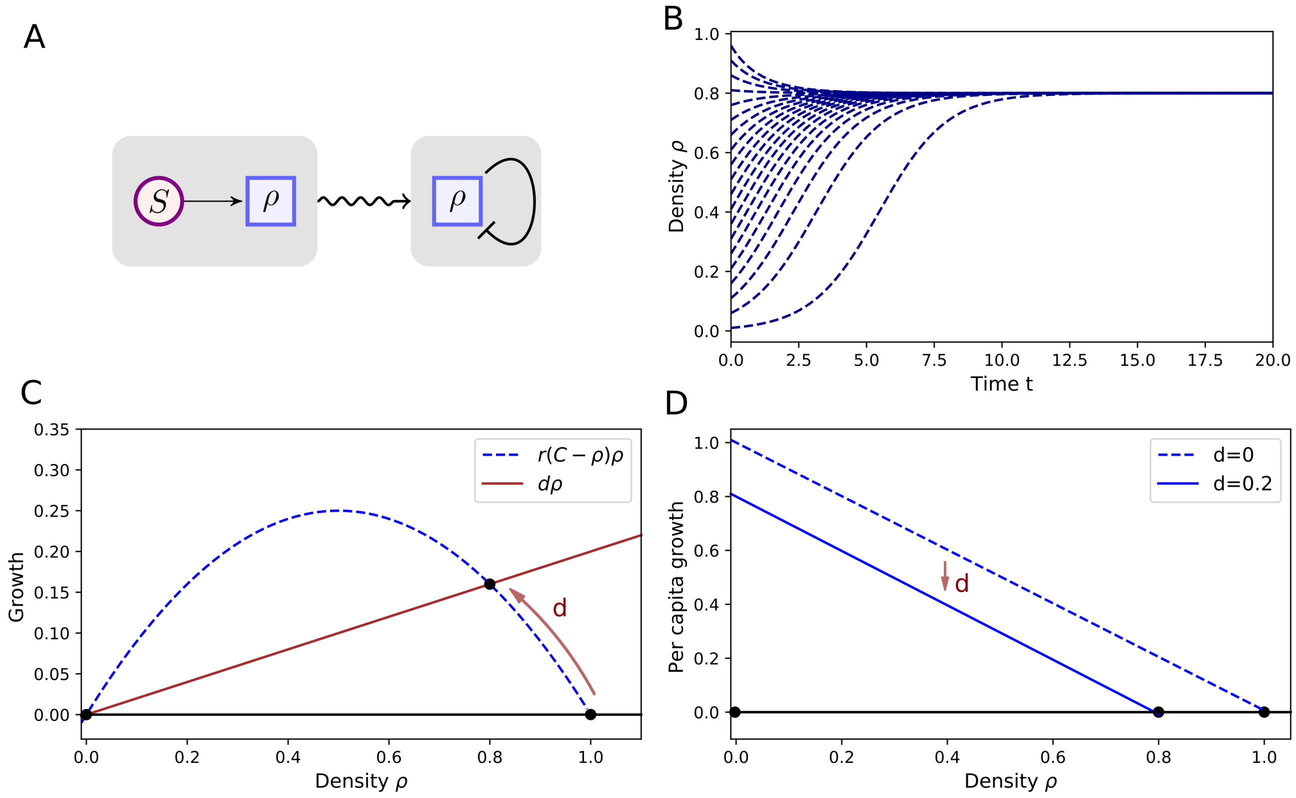


**Figure S1.** Consumption of a nutrient (concentration $S$) by a microbial species (density $\rho$) can be described by the logistic growth equation (S3). (A) Resource-dependent growth equations are reduced (wavy arrow) to a single logistic equation. The availability of the substrate limits the growth. (B) Regardless of its initial value, the density will eventually approach the equilibrium state, determined by the carrying capacity. (C) The dilution rate reduces the growth of a species. The equilibrium state is determined by the intersection of the growth term and the dilution term. (D) Equivalent representation as in (C), showing that the per capita growth is reduced by the dilution rate $d$. As a result of the introduction of the dilution rate, the carrying capacity is reduced. Parameter values are given in Table S3

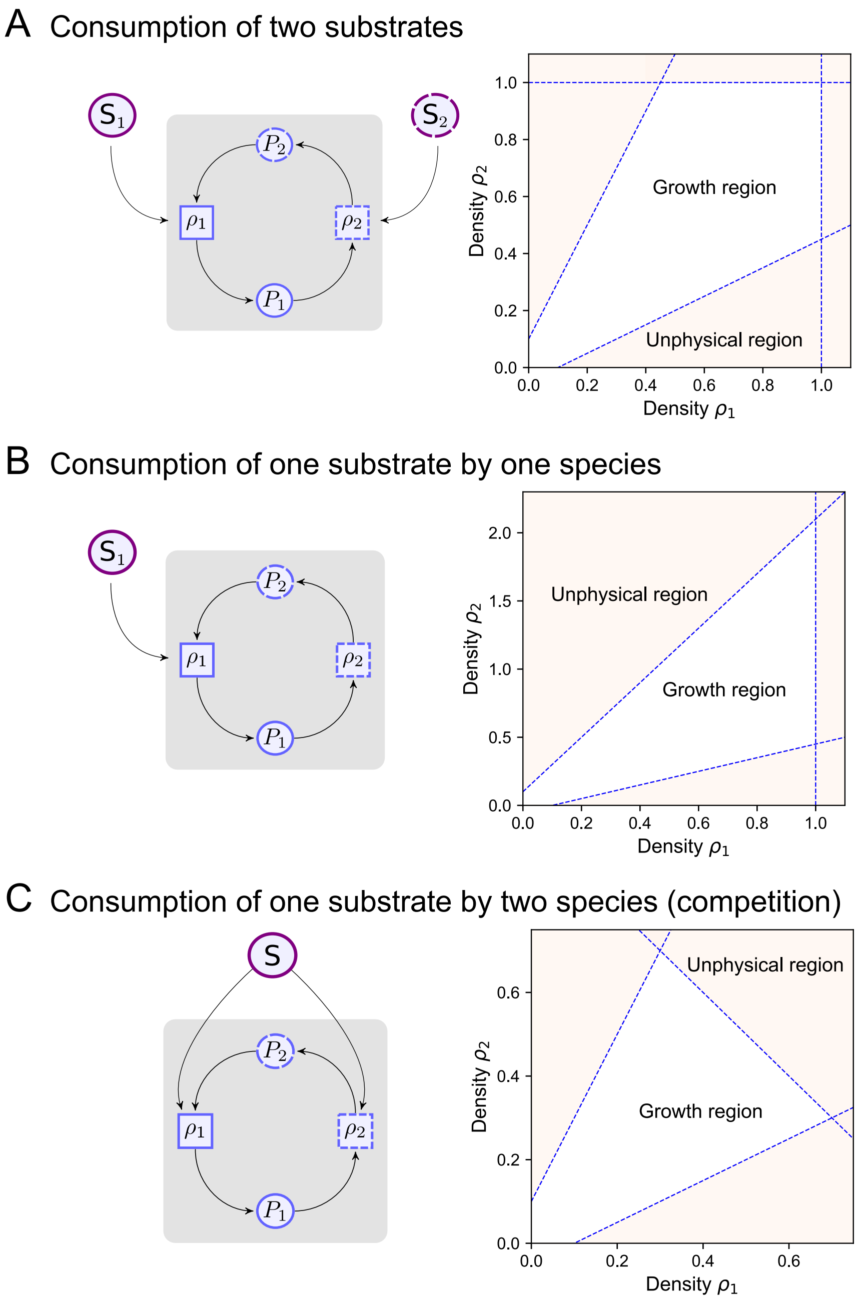


**Figure S2.** Growth limitation of both species by the substrates and corresponding physical growth region in the phase plane. (A) Mutualistic cross-feeding (via cross-fed nutrients with concentrations $P_1$ and $P_2$) with additional consumption of two substrates with concentrations $S_1$ and $S_2$, by species 1 and 2, respectively. (B) Mutualistic cross-feeding with growth limitation of one species by a substrate. (C) Mutualistic cross-feeding with competition for the same substrate.

### S2 Reduction of nutrient-explicit mutualistic cross-feeding equations

Mutual cross-feeding between two species (densities $\rho_1$ and $\rho_2$) via nutrients with concentrations $P_1$ and $P_2$ is modeled with similar consumer-resource equations as for one species. However, an external nutrient (concentration $S$) is required to avoid unphysical divergent growth, which we will refer to as the substrate to clearly distinguish it from the cross-feeding nutrients. The substrate can be seen as the source of energy for the mutualistic species to grow and to produce the cross-feeding nutrients.

We can distinguish the following different mutualistic systems:

- Consumption of two different substrates by the two species (Fig S2 A)
- Consumption of a substrate by only one species (Fig S2 B)
- Competition for a substrate by the two species (Fig S2 C)

Each of these systems has different boundaries for the physical growth region but the qualitative dynamics remains the same as the mutualistic cross-feeding is not affected by the substrate. In the following, we first summarize how these systems are described by chemostat equations. Then, we explain how resource-explicit equations of these different motifs are reduced. We first focus on the system with two different substrates. Afterwards, we make the connection to the other two situations and show the similarities.
Note that the reduction is done in a similar fashion for batch and for chemostat. The only difference is that in batch the dilution rate $d$ is set to $d = 0$.

#### Overview of the different explicit chemostat equations

The different parameters are defined in Table S5.

- Mutualistic cross-feeding between two species (densities $\rho_1$ and $\rho_2$) via the cross-fed nutrients with concentrations $P_1$ and $P_2$, with consumption of two different substrates with concentrations $S_1$ and $S_2$ (Fig S2A):

$$\begin{aligned}
\frac{d\rho_1}{dt} &= F_1(S_1,P_1)\rho_1 - d\rho_1,\\
\frac{d\rho_2}{dt} &= F_2(S_2,P_2)\rho_2 - d\rho_2,\\
\frac{dS_1}{dt} &= d\tilde{S_1} - dS_1 - \nu_{s1}F_1(S_1,P_1)\rho_1,\\
\frac{dS_2}{dt} &= d\tilde{S_2} - dS_2 - \nu_{s2}F_2(S_2,P_2)\rho_2,\\
\frac{dP_1}{dt} &= d\tilde{P_1} - dP_1 - \nu_{p1}F_1(S_1,P_1)\rho_1 + a_{12}F_2(S_2,P_2)\rho_2,\\
\frac{dP_2}{dt} &= d\tilde{P_2} - dP_2 - \nu_{p2}F_2(S_2,P_2)\rho_2 + a_{21}F_1(S_1,P_1)\rho_1.
\end{aligned} \tag{S7}$$

- Mutualistic cross-feeding between two species, with consumption of one substrate (concentration $S_1$) by one species (Fig S2B):

$$\begin{aligned}
\frac{d\rho_1}{dt} &= F_1(S_1,P_1)\rho_1 - d\rho_1,\\
\frac{d\rho_2}{dt} &= F_2(P_2)\rho_2 - d\rho_2,\\
\frac{dS_1}{dt} &= d\tilde{S_1} - dS_1 - \nu_{s1}F_1(S_1,P_1)\rho_1,\\
\frac{dP_1}{dt} &= d\tilde{P_1} - dP_1 - \nu_{p1}F_1(S_1,P_1)\rho_1 + a_{12}F_2(P_2)\rho_2,\\
\frac{dP_2}{dt} &= d\tilde{P_2} - dP_2 - \nu_{p2}F_2(P_2)\rho_2 + a_{21}F_1(S_1,P_1)\rho_1.
\end{aligned} \tag{S8}$$

- Mutualistic cross-feeding between two species, with competition for a substrate with concentration $S$ (Fig S2C):

$$\begin{aligned}
\frac{d\rho_1}{dt} &= F_1(S,P_1)\rho_1 - d\rho_1,\\
\frac{d\rho_2}{dt} &= F_2(S,P_2)\rho_2 - d\rho_2,\\
\frac{dS}{dt} &= d\tilde{S} - dS - \nu_{s1}F_1(S,P_1)\rho_1 - \nu_{s2}F_2(S,P_2)\rho_2,\\
\frac{dP_1}{dt} &= d\tilde{P}_1 - dP_1 - \nu_{p1}F_1(S,P_1)\rho_1 + a12F_2(S,P_2)\rho_2,\\
\frac{dP_2}{dt} &= d\tilde{P}_2 - dP_2 - \nu_{p2}F_2(S,P_2)\rho_2 + a_{21}F_1(S,P_1)\rho_1.
\end{aligned} \tag{S9}$$

#### Reduction

We reduce Eqs (S7) for the case of two different substrates, after which we discuss the differences for the two other cases. Essentially, the reduction consists of two steps: a linearization of the growth rate via a Taylor approximation and applying conservation of biomass when $t \gg 1/d$.

##### *First simplification: Taylor approximation for limiting nutrient concentrations*

The growth rates $F_i(S_i,P_i)$, with $i=1,2$ are assumed to depend obligatory on $S_i$ and $P_i$. This has the following implications for the growth rate $F_i(S_i,P_i)$:

$$\begin{aligned}
F_i(0,0) &= 0\\
\tfrac{\partial F_i}{\partial S_i}(0,0) &= 0\\
\tfrac{\partial F_i}{\partial P_i}(0,0) &= 0
\end{aligned}$$

In fact, all partial derivatives to one of the variables $S_i$ or $P_i$ are zero, otherwise there is a net growth if only one of the nutrients is missing.

The first non-zero term of the Taylor approximation for the growth rate $F_i(S_i,P_i)$, for limiting nutrient concentrations, is then:

$$F_i(S_i,P_i) \approx \frac{\partial^2 F_i}{\partial S_i \partial P_i}(0,0)S_iP_i$$

An example for which these conditions are fulfilled are Monod-like growth rates:

$$F_i(S_i,P_i) = \mu_i \frac{S_i}{S_i+K_{Si}}\frac{P_i}{P_i+K_{Pi}} \tag{S10}$$

With $\mu_i$ the maximal growth rate and $K_{Si}$ and $K_{Pi}$ the half-saturation constants. At low concentrations $S_i$ and $P_i$, this leads to:

$$F_i(S_i,P_i) \approx \frac{\mu_i}{K_{Si}K_{Pi}}S_iP_i$$

##### *Second simplification: conservation of biomass for $t \gg 1/d$*

As for the case for the growth of a single species, we can linearly combine the equations to obtain the following conservation laws for $t \gg 1/d$:

$$\begin{aligned}
S_1 - \nu_{s1}\rho_1 &= \tilde{S}_1,\\
S_2 - \nu_{s2}\rho_2 &= \tilde{S}_2,\\
P_1 + a_1\rho_1 - \nu_2\rho_2 &= \tilde{P}_1,\\
P_2 + a_2\rho_2 - \nu_1\rho_1 &= \tilde{P}_2.
\end{aligned} \tag{S11}$$

This description is limited to positive values of the nutrients. Hence, the physical phase space is limited to:

$$\begin{aligned} &C_{s1} - \rho_1 > 0, \\ &C_{s2} - \rho_2 > 0, \\ &C_{p1} + b_1\rho_2 - \rho_1 > 0, \\ &C_{p2} + b_2\rho_1 - \rho_2 > 0. \end{aligned} \tag{S12}$$

These correspond to the nullclines in the case that $d = 0$ (growth in batch) as well are these the asymptotes of the hyperbolic nullclines when $d \neq 0$. These equations form the physical growth region in the phase plane (Fig S2 A and B).

##### ***Elimination of the equations for the substrates and reduction to two variables***

Performing the reduction on the system with two substrates, Eq (S7), the following equations for $\rho_1$ and $\rho_2$ are obtained:

$$\begin{aligned} \frac{d\rho_1}{dt} &= r_1(C_{s1} - \rho_1)(C_{p1} + b_1\rho_2 - \rho_1)\rho_1 - d\rho_1, \\ \frac{d\rho_2}{dt} &= r_2(C_{s2} - \rho_2)(C_{p2} + b_2\rho_1 - \rho_2)\rho_2 - d\rho_2. \end{aligned} \tag{S13}$$

The growth rates $r_1$, $r_2$, carrying capacities $C_{s1}$, $C_{s1}$, $C_{p1}$, $C_{p2}$ and production constants $b_1$, $b_2$ are defined as in table S6.

The same reduction method can be applied to the system with one substrate for one species (Eq (S8)) and the system with competing species (Eq S9), resulting in the following equations:

- Mutualistic cross-feeding between two species, with consumption of one substrate by one species (Fig S2B):

$$\begin{aligned} \frac{d\rho_1}{dt} &= r_1(C_s - \rho_1)(C_{p1} + b_1\rho_2 - \rho_1)\rho_1 - d\rho_1, \\ \frac{d\rho_2}{dt} &= r_2(C_{p2} + b_2\rho_1 - \rho_2)\rho_2 - d\rho_2. \end{aligned} \tag{S14}$$

- Mutualistic cross-feeding between two species, with competition for one substrate (Fig S2C):

$$\begin{aligned} \frac{d\rho_1}{dt} &= r_1(C_s - c\rho_2 - \rho_1)(C_{p1} + b_1\rho_2 - \rho_1)\rho_1 - d\rho_1, \\ \frac{d\rho_2}{dt} &= r_2(C_s - c\rho_2 - \rho_1)(C_{p2} + b_2\rho_1 - \rho_2)\rho_2 - d\rho_2. \end{aligned} \tag{S15}$$

  The parameter $c$ is defined as $c = \frac{\nu_{s2}}{\nu_{s1}}$ (table S6). It is the relative consumption of the substrate by species $\rho_2$ to species $\rho_1$.

**Table S4.** Variables for the two-species system

| Variables | Biological meaning |
|---|---|
| $\rho$ | microbial density |
| $\rho_1$ | species 1 |
| $\rho_2$ | species 2 |
| $S_1$ | substrate for species 1 |
| $S_2$ | substrate for species 2 |
| $P_1$ | cross-feeding nutrient consumed by species 1 |
| $P_2$ | cross-feeding nutrient consumed by species 2 |

**Table S5.** Parameters of the chemostat model for 2 mutualists with 2 substrates ($i,j = 1,2$ and $i \neq j$).

| Parameters | Biological meaning | Value |
|---|---|---|
| $d$ | dilution rate | 0.2 |
| $\tilde{S}_i$ | inflow concentration of substrate $i$ | 1 |
| $\tilde{P}_i$ | inflow concentration of cross-fed nutrient $i$ | 0.1 |
| $F_i(S_i,P_i)$ | Growth rate of species $i$, function of $S_i$ and $P_j$. | $\mu_i \frac{S_i}{K_{si}+S_i} \frac{P_j}{K_{pi}+P_j}$ |
| $\mu_i$ | maximal growth rate of species $i$ | 2 |
| $K_{si}$ | half-saturation constant of species $i$ for $S_i$ | 2 |
| $K_{pi}$ | half-saturation constant of species $i$ for $P_i$ | 1 |
| $\nu_{si}$ | Consumption rate of $S_i$ by species $i$ | 1 |
| $\nu_{pi}$ | Consumption rate of $P_i$ by species $i$ | 1 |
| $a_{ij}$ | production rate of $P_i$ by species $j$ | 2 |

**Table S6.** Parameters of the reduced model of 2 mutualists ($i,j = 1,2$ and $i \neq j$). In case of symmetric parameters, the subscripts $i$ and $j$ are removed.

| Parameters | Biological meaning | Relation to chemostat | Value |
|---|---|---|---|
| $d$ | dilution rate | $d$ | 0.2 |
| $r_i$ | growth rate of species $i$ | $\frac{\partial^2 F_i}{\partial S_i \partial P_i}(0,0)\nu_{si}\nu_{pi}$ | 1 |
| $C_{si}$ | Carrying capacity: growth limitation by the substrate $i$ | $\frac{\tilde{S}_i}{\nu_{si}}$ | 1 |
| $C_{pi}$ | Growth limitation of species $i$ by cross-fed nutrient $j$ | $\frac{\tilde{P}_j}{\nu_i}$ | 0.1 |
| $b_i$ | Growth stimulation through cross-feeding (via $P_i$) | $\frac{a_{ij}}{\nu_{pi}}$ | 2 |
| $c$ | Competitive strength (in case of competition for $S$) | $\frac{\nu_{sj}}{\nu_{si}}$ | 1 |

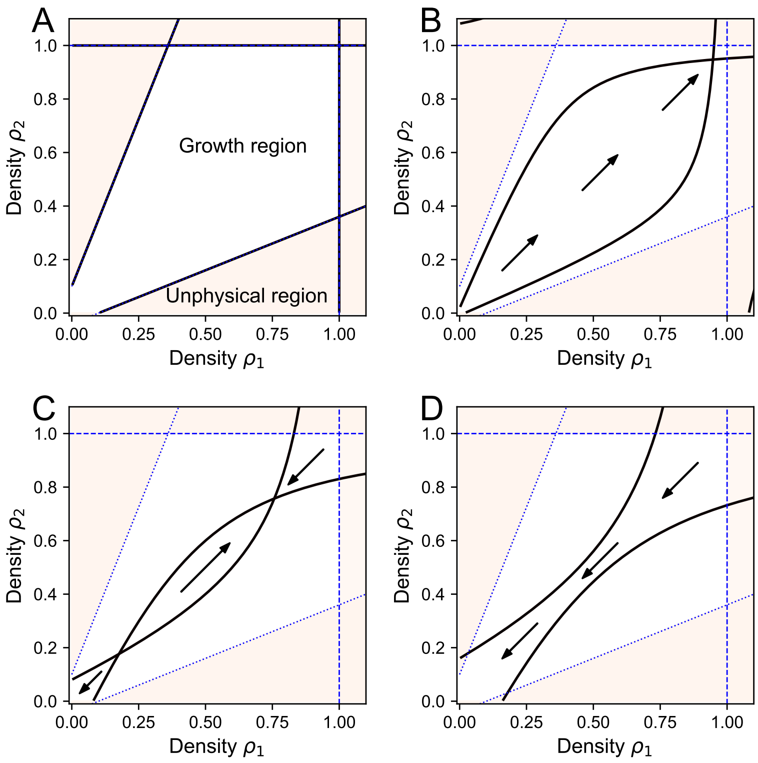


**Figure S3.** Phase plane of the mutualistic system with 2 substrates for different values of the dilution rate $d$. (A) dynamics in batch ($d = 0$). The nullclines are the same as the limiting functions, Eq.(S12). (B) Coexistence at low dilution ($d = 0.08$). The nullclines become hyperbola with the asymptotes determined by the limiting functions of the growth region. (C) Bistability at intermediate dilution ($d = 0.3$). Bistability arises as the nullclines intersect twice, thereby creating a saddle point as the separation between extinction and coexistence. (D) Extinction at large dilution ($d = 0.5$). There is no survival as the only fixed point is the state where the species are extinct.

**Similarities between the different systems**

The three different mutualistic systems behave similarly: the mutualistic cross-feeding is described in the same way in all equations, but the dependence on the substrate differs. Growth limitation via one substrate (Eq (S8)) can be seen to be a special case of the limitation with two substrates (Eq (S7)), when the consumption of the second substrate is negligible (see Fig S2 A and B).

The general mutualistic system (Eq (S7)) is analysed in the phase plane for different values of the dilution rate, by considering the nullclines determined by $d\rho_1/dt = 0$ and $d\rho_2/dt = 0$ (Fig S3). The growth region is limited by Eqs (S12). The consumption of the substrates determines the growth limitation at high densities, the growth region being limited by $\rho_1 = C_{s1}$ and $\rho_2 = C_{s2}$. As described in the main text, the behavior changes from global survival to bistability to global extinction with increasing values of $d$.

Competition for a substrate introduces an extra interdependence of the two species as the growth is limited by the substrate, which decreases faster due to the presence of the other species. In contrast to the case of consumption of two separate substrates, the feasible region at high densities being limited by $c\rho_2 + \rho_1 = C_s$. This means that the total carrying capacity depends on the density of the other species. The dynamics of this system remains qualitatively the same as the nullclines are similar hyperbolic functions, leading to coexistence at low dilution, bistability at intermediate dilution and extinction at high dilution (Fig S4). As the carrying capacity is reduced by the other species, the densities at the coexisting state is smaller than in the case of pure mutualism.

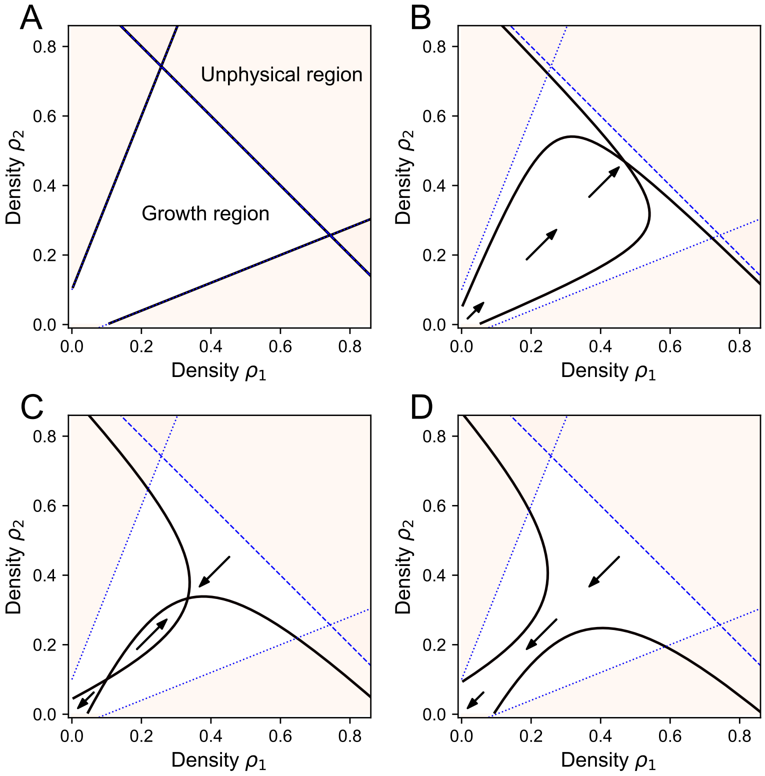


**Figure S4.** Phase planes of the mutualistic system with competition for different values of the dilution rate $d$. (A) dynamics in batch ($d = 0$): the nullclines are the limiting functions of the growth region. (B) Coexistence at low dilution ($d = 0.05$). The nullclines become hyperbola with the asymptotes determined by the limiting functions of the growth region.(C) Bistability at intermediate dilution ($d = 0.2$). Bistability arises as the nullclines intersect twice, thereby creating a saddle point as the separation between extinction and coexistence. (D) Extinction at large dilution ($d = 0.3$). There is no survival as the only fixed point is the state where the species are extinct.

### S3 Analysis for asymmetric parameter values

In the main text, we analyzed the mutualistic system using symmetric parameters, so that the following growth equation with an Allee effect is obtained:

$$\frac{d\rho}{dt} = r(C-\rho)(\rho+a)\rho - d\rho \tag{S16}$$

**Table S7.** Parameters for growth with Allee effect (for symmetric parameters)

| Parameters | Biological meaning | Calculation | Value |
|---|---|---|---|
| $d$ | dilution rate | $d$ | 0.2 |
| $a$ | Allee effect | $\frac{C_p}{b-1}$ | 0.1 |
| $r_a$ | growth rate | $r(b-1)$ | 1 |
| $C_a$ | carrying capacity | $C_s$ | 1 |

Here, we show how the dynamics can be analyzed in the case of different parameter values for the two species. To do this, we assume that $C_{p1} \ll C_{s1}$ and $C_{p2} \ll C_{s2}$, meaning that the inflow of cross-feeding nutrients is small in comparison with the inflow of substrate.

As we are interested in the Allee effect which is the increased fitness at low densities, we approximate the nullclines of Eq (S13) at $\rho_i \ll C_{si}$ $(i = 1,2)$ by:

$$\begin{aligned} \frac{d\rho_1}{dt} &= C_{p1} + b_1\rho_2 - \rho_1 - \frac{d}{C_{s1}r_1} = 0, \\ \frac{d\rho_2}{dt} &= C_{p2} + b_2\rho_1 - \rho_2 - \frac{d}{C_{s2}r_2} = 0. \end{aligned} \tag{S17}$$

For bistability to exist, these nullclines need to intersect in the positive quadrant of the phase plane. The intersection point is determined by the following densities of $\rho_1$ and $\rho_2$:

$$\begin{aligned} \rho_1^s &= \frac{-(b_1C'_{p2} + C'_{p1})}{b_1b_2 - 1}, \\ \rho_2^s &= \frac{-(b_2C'_{p1} + C'_{p2})}{b_1b_2 - 1}, \end{aligned} \tag{S18}$$

where we defined the following parameters $(i = 1,2)$:

$$C'_{pi} = C_{pi} - \frac{d}{C_{si}r_i}. \tag{S19}$$

Here, $C'_{p1}$ is the solution of the nullcline $d\rho_1/dt = 0$ when $\rho_2 = 0$. As a result, $C'_{p1}$ is the equilibrium density of $\rho_1$ when $\rho_2 = 0$. Similarly, $C'_{p2}$ is the equilibrium density of $\rho_2$ when $\rho_1 = 0$.

When $b_1b_2 < 1$, the dynamics correspond to logistic growth, as the growth is limited by inflow of the cross-feeding nutrients. Bistability thus only exists for $b_1b_2 > 1$, as this allows the shape of the physical growth region to be such that the nullclines can intersect twice (Fig S5A, C). The values of $C'_{p1}$ and $C'_{p2}$ determine the position of the saddle node in the phase plane.

We thus obtain the two following necessary conditions for bistability:

- $b_1b_2 > 1$
- $b_1C'_{p2} + C'_{p1} < 0$ and $b_2C'_{p1} + C'_{p2} < 0$

As long as the above conditions for bistability are fulfilled, then depending on the sign of $C'_{p1}$ and $C'_{p2}$ it is possible to find bistability between coexistence of both species and the survival of a single species (density $C'_{pi}$), see Fig S5D. The same qualitative reasoning as in the main text holds, where we restricted our analysis to bistability between coexistence and extinction of both species.

The following types of bistability are thus distinguished for asymmetric parameters:

- Type I: $C'_{p1} < 0$ and $C'_{p2} < 0$, bistability between coexistence and extinction of both species
- Type II: $C'_{p1} > 0$ and $C'_{p2} < 0$, bistability between coexistence and survival of species 1
- Type III: $C'_{p1} < 0$ and $C'_{p2} > 0$, bistability between coexistence and survival of species 2

This is summarized in Fig S5B, where we show region with different types of dynamics (monostability or bistability) as a function of $C'_{p1}$ and $C'_{p2}$. Both parameters decrease with the dilution rate, so that it is possible to follow a trajectory (1 and 2 in Fig S5B) to describe the effect of the dilution. Dilution thus turns monostability (coexistence of 2 species) into bistability. Eventually, if the dilution becomes too large, bistability is lost when the saddle point touches the stable equilibrium of coexistence in a saddle-node bifurcation. This results in the extinction of both species for all initial densities (monostability).

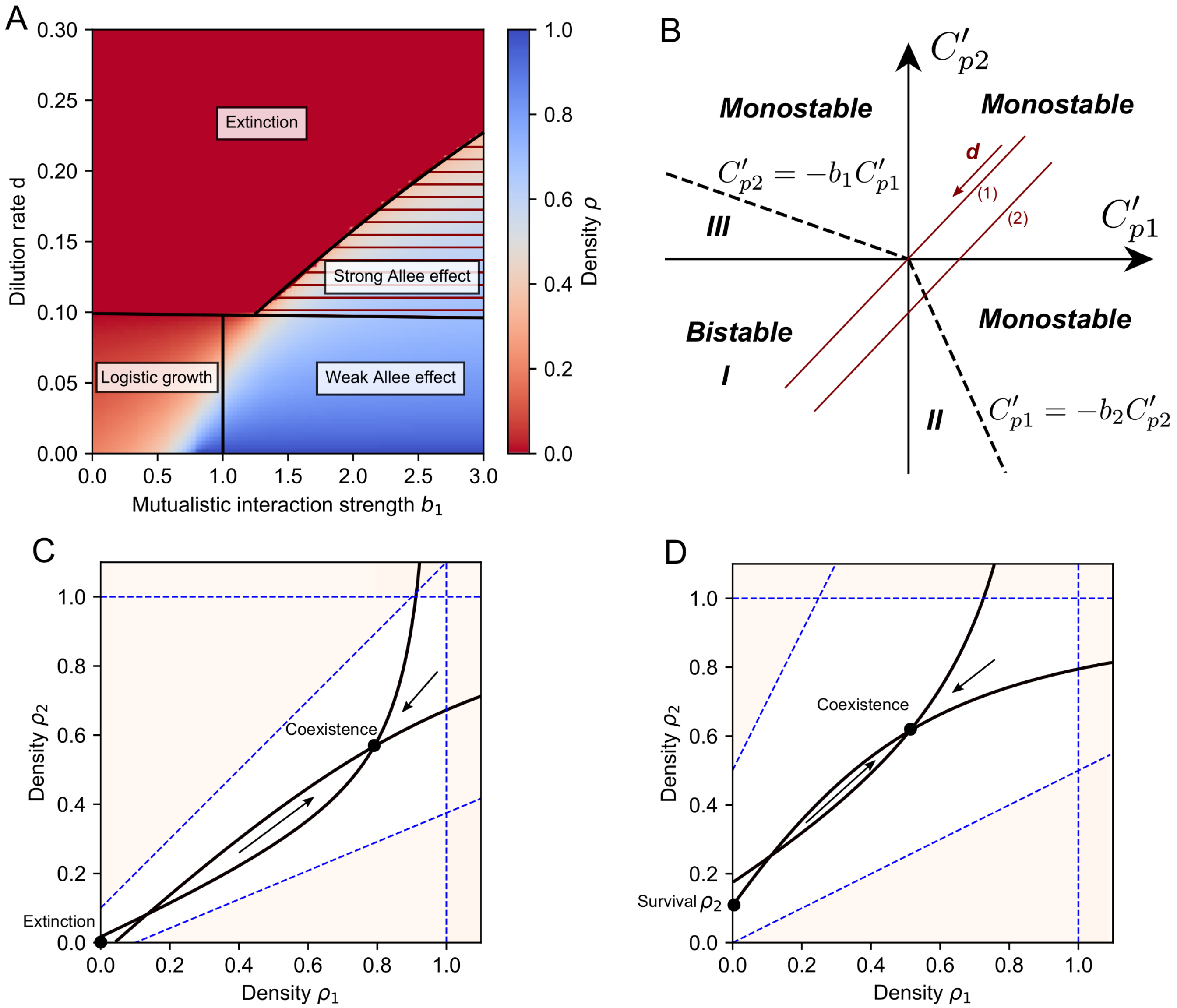


**Figure S5.** The effect of asymmetric parameters. (A) Influence of the asymmetry of the mutualistic interaction strength. Parameter $b_2 = 1$ was fixed, while $b_1$ varies showing the necessary condition $b_1 b_2 > 1$ for bistability. (B) The introduced parameters $C'_{p1}$ and $C'_{p2}$ (Eq (S19)) predict the possiblity of bistability as these determine the position of the saddle node, given by Eq (S18). The system exhibits bistability type I (coexistence or extinction) for $C'_{pi} < 0$ $(i = 1, 2)$, type II (coexistence or survival species 1) for $C'_{p2} < 0$ and $0 < C'_{p1} < -b_2 C'_{p2}$ and type III (coexistence or survival species 2) for $C'_{p2} < 0$ and $0 < C'_{p1} < -b_2 C'_{p2}$. Decreasing the dilution will decrease $C'_{p1}$ and $C'_{p2}$ so that the system becomes bistable, trajectories are shown for $C_{p1} = C_{p2}$ (1) and $C_{p1} > C_{p2}$ (2). When the dilution becomes too large there is extinction of the species. (C) Phase plane showing bistability type I for asymmetric values of $b$ $(b_1 = 2.4,\ b_2 = 1)$ . (D) Phase plane showing bistability type III for $C_{p1} = 0$ and $C_{p2} = 0.5$.
